# Diminished social memory and hippocampal correlates of social interactions in chronic social defeat stress susceptibility

**DOI:** 10.1101/2024.08.06.606823

**Authors:** Amanda Larosa, Tian Rui Zhang, Alice S. Wong, Y. H. Fung Cyrus, Xiong Ling Yun (Jenny) Long, Benjamin C. M. Fung, Tak Pan Wong

## Abstract

**Background:** The susceptibility to chronic stress has been associated with depression, a mood disorder which highly implicates the hippocampus. Hippocampal contribution to stress susceptibility has been supported by findings in mice following chronic social defeat stress (CSDS). However, little is known of the role of hippocampal activity in determining the development of stress susceptibility.

**Methods:** We used the UCLA miniscope to longitudinally measure the activity of dorsal CA1 hippocampal neurons across CSDS. Apart from examining the representation of social information by these neurons, we also compared social memory in mice that were susceptible or resilient to CSDS.

**Results:** We observed more stable dCA1 correlates of social interaction and social memory in CSDS resilience. Such changes were absent in CSDS susceptible mice and accompanied by greater social memory impairments.

**Conclusions:** CSDS susceptibility may be supported by hippocampal social cognitive processes, reflected in diminished hippocampal representations of social information and a greater impairment in social memory.

## Introduction

Chronic stress represents a significant risk factor in the pathogenesis of depression, a debilitating mood disorder impacting an estimated 280 million individuals globally (*1, 2*). The hippocampus is highly examined in neurobiological investigations of depression (*3*). Differential hippocampal changes in stress-exposed groups that go on to develop depression-like features, termed susceptible, compared to those that appear resilient to stress have been identified. Hippocampal hyperactivity with stress susceptibility is among the observed changes in both depressed patients responding to negative stimuli and mouse models of depression (*4–6*). Using a social stressor called chronic social defeat stress (CSDS) to separate mice into susceptible and resilient populations (*7*), we found more hippocampal CA1 neurons were reactivated by the CSDS experience in susceptible mice (*8*). Changes in hippocampal activity invariably led to alterations in its function, which could be rooted in social cognition.

The hippocampus supports social cognition. In humans, hippocampal activity during facial recognition correlates with how well participants report knowing a presented face (*9*). Findings in rodents support crucial roles of hippocampal subregions, including the dorsal CA2, ventral CA3 and CA1 (*10–12*), in social memory. Emerging findings also suggest that the dorsal CA1 (dCA1) contributes to social information processing. dCA1 neurons preferentially fire according to the location of a conspecific (*13, 14*), which is likely crucial for informing the decision to engage in prosocial (e.g., kin approach) or antisocial behaviours (e.g., avoidance). The dCA1 is also essential for the recognition of social identity (*15, 16*). Whether changes in dCA1 representation of social targets are related to the susceptibility to a social stressor remains unclear.

To examine dCA1 representation of social information during CSDS, we longitudinally performed *in vivo* calcium imaging of dCA1 neurons throughout CSDS in male C57BL/6 mice. We hypothesize that the altered hippocampal activity in stress susceptible mice may impact the processing of social information by the dCA1 and social memory.

## Methods and Materials

### Animals

Male adult C57BL/6 (2-3-month-old) and CD1 mice (>6-month-old) were obtained from *Charles River*. All experiments were approved by the Facility Animal Care Committee and followed the guidelines from the Canadian Council on Animal Care (protocol#.: DOUG-5935).

### Surgeries

Under isoflurane, 253 nL of AAV2/9-SYN-GCaMP6f virus at 23 nL/sec was stereotaxically injected in the dCA1 (bregma: AP:-1.60; ML:+1.70; DV:-1.40 or AP:-1.90; ML:+1.40; DV:-1.10, both coordinates achieved comparable results). One week later, a gradient refractive index (GRIN) lens (pitch:0.25, diameter:1.8 mm, length:4 mm, *Edmund Optics*) was implanted above the injection site, with cortical tissue above it aspirated. The GRIN lens was fixed in position using cyanoacrylate and dental cement. Three weeks later, a baseplate for attaching the miniscope was cemented on the head surface.

### Chronic Social Defeat Stress (CSDS)

C57BL/6 mice were attacked by CD1 aggressors during social defeat as described (*8*). CSDS consisted of 8 episodes of defeat. Mice were attacked by up to 12 attacks in a maximum period of 5 minutes in each defeat. Following defeat, C57BL/6 mice were housed with the CD1 aggressor for 24 hours. A perforated partition separated defeated mice from aggressors during cohousing to allow for the presence of sensory stressors. A new aggressor was used in each defeat episode to prevent habituation from the same aggressor. Control non-stressed C57BL/6 mice were handled and pair-housed in neighbouring partitions for 8 days. After CSDS or pair-housing, social behaviors of stressed and control mice were examined in a social interaction test (SIT, (*17*)).

The SIT consisted of two 150-second sessions in an open field (44 cm x 44 cm). An empty perforated enclosure (10 cm x 5 cm x 30 cm) was placed in the center of the north side of the open field during session 1. During session 2, a novel CD1 mouse was placed in the enclosure. Both sessions were performed under ambient red light and static white noise (60 dB). Time spent in the interaction zone (10 cm around the enclosure) during SIT was estimated using TopScan LITE (*Clever system Inc.*). SI ratio equals to interaction time-session 2 (CD1) / interaction time-session 1 (empty). Stressed mice with SI ratio > 1 are considered as resilient. All other stressed mice are susceptible mice. GRIN lens placement was histologically confirmed after the SIT.

### Social memory tests

On day 1, subject mice were placed in an open field for two 8-minute sessions separated by 2 hours. In the first session, the open field contained an empty wire cup and a novel CD1 A-containing wire cup, while in the second session the wire cups contained the now familiar CD1 A and novel CD1 B. On day 2, subject mice were reintroduced to the open field which contained familiar CD1 A and an empty wire cup. All sessions were performed under ambient red light and static white noise. Positions of the wire cups were counterbalanced.

### *In vivo* calcium imaging

Using the UCLA Miniscope (v3), we conducted *in vivo* calcium imaging of dCA1 neurons during cohousing with a conspecific or an aggressor after the 2^nd^, 5^th^ and 8^th^ episode of social defeat and SIT. Mice were separated from the cohoused control or aggressor by a perforated partition, with the cage cover removed to allow cable access and mouse behavior recording.

Miniscope recording was performed via the Miniscope-DAQ-QT-Software (Aharoni-Lab, *github*). These recordings were concatenated, motion-corrected and aligned using NoRMCorre (*18*). Calcium signals from each cell segment were identified and extracted using CNMF-E (*19*).

Overlapping segments (>60%) were discarded. The rising phase of these signals was binarized for estimating inferred spikes. Mouse body parts were annotated using *DeepLabCut* (***20***) to generate head position and heading direction vectors. All calcium and behavioral data were down sampled to 5 Hz with customized MATLAB codes for additional analyses.

### Social ensembles

dCA1 neurons were sorted according to their activity during bouts of social interaction. Relative to the head of social target, social interaction bouts were defined as <10 cm between the head of the subject mouse, and <50° from the heading direction of the subject mouse. Only social interactions lasting for at least 5 frames (1 second) were used. We identified neurons that were active during at least 40% of the social interaction bouts and calculated the mean activity of these neurons to construct the social ensemble vector. We found that the cosine similarity index (CSI, (*21, 22*)), which was calculated from the ensemble activity vector (***C***) and the social interaction bout vector (binary logical values of social interaction frames ***B***; CSI ***=* 2*B*** · ***C***/(|***B***|^**2**^ · |***C***|^**2**^)), of social ensemble vector is significantly higher than 95% of shuffled social interaction bout vectors (10,000 shuffled vectors). Social ensemble vector also showed poorer similarity with other behavioral vectors such as subject mouse running speed and head direction.

Ensemble size was calculated by dividing the number of dCA1 neurons within a social ensemble by the total number of registered neurons within the same recording. Neuron reactivation was tracked using *CellReg* (*23*). Spatial information of dCA1 neurons was estimated as previously described (*24*) using spatial bins of 4 cm^2^.

### Statistical analysis

All statistical analyses were performed using *JMP Pro 15* and *GraphPad Prism 10.* Normality of data was examined by the Shapiro-Wilk’s test. Two-way ANOVA and post hoc Tukey test were used to analyze calcium imaging data. For behavior measures, one-way ANOVA and post hoc Tukey test were used. Kruskal–Wallis test and post hoc Wilcoxon tests were used for data that are not normally distributed. Student’s t-test and Mann-Whitney test were used for parametric and non-parametric comparisons. All data were presented as mean ± SEM.

## Results

### Mouse behaviors during CSDS cohousing and SIT

We longitudinally monitored dCA1 activity in conjunction with social behavior using the miniscope to determine how neuronal correlates of social interactions evolved throughout CSDS. *In vivo* calcium imaging was performed during the cohousing that followed a defeat episode where both the C57BL/6 subject and CD1 aggressor (or non-stressed control) were freely moving, or during the social interaction test (SIT) on day 9 (**Fig. 1a**). To examine the long-term effect of CSDS on hippocampal activity, *in vivo* calcium imaging was performed 1 hour post-defeat. CSDS susceptibility and resilience were determined from behavioral performance in the SIT (See methods). We identified 11 resilient and 8 susceptible mice which were compared with 9 non-stressed controls. Briefly, in the SIT control and resilient mice spent more time in the area surrounding a CD1 mouse-containing enclosure, termed the interaction zone, than susceptible mice **(fig. S1a, b)**. Susceptible mice instead spent more time in the corner zones (**fig. S1c**). No differences in weight change, or the latency to first attack and attack number during CSDS were observed between the two stressed groups **(fig. S1d)**.

**Figure 1:**
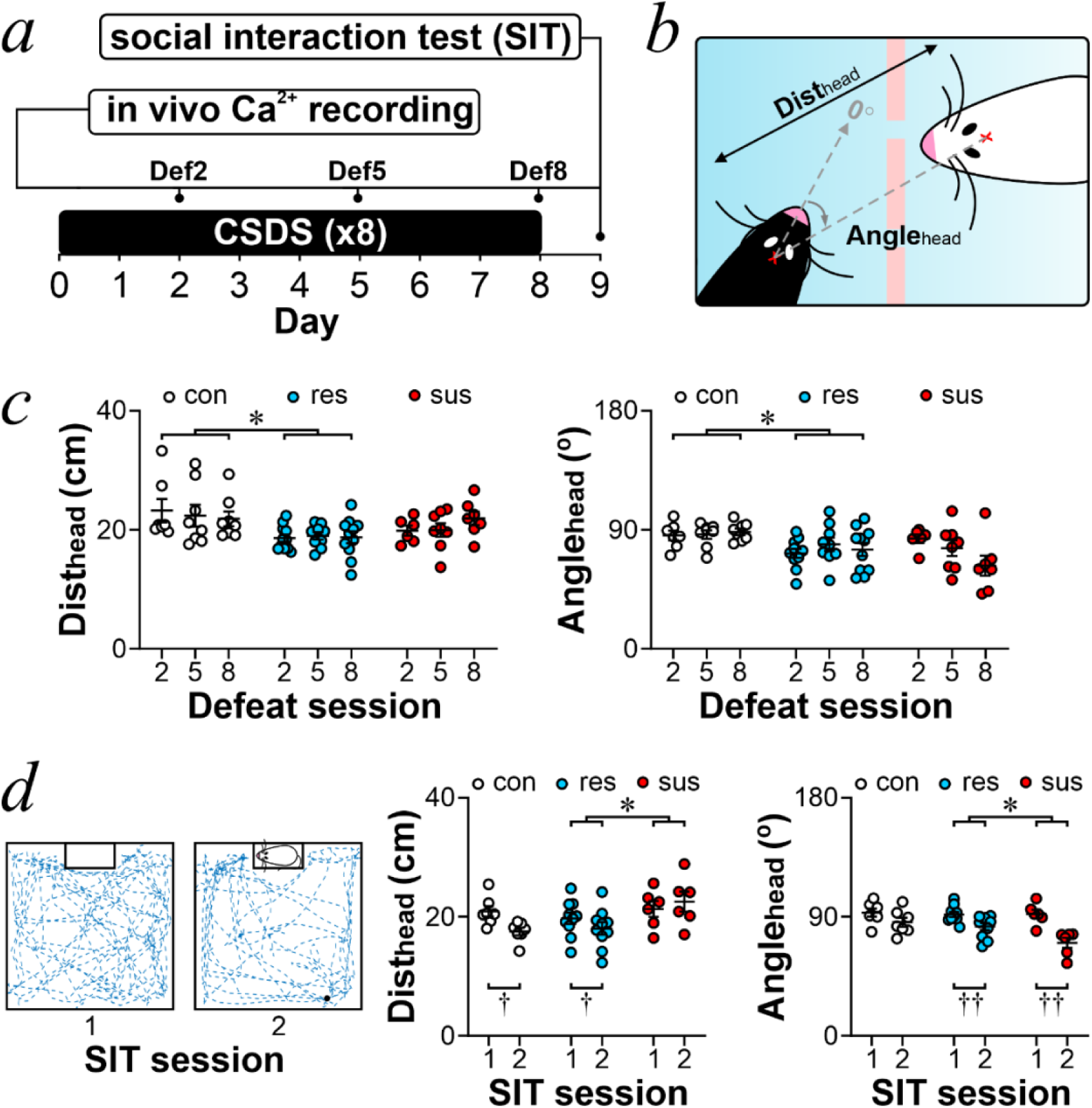
Differences in behavior between susceptible and resilient mice during cohousing post-CSDS and SIT. (**a**) Schematic of experiment timeline. On day 1, chronic social defeat stress (CSDS) began and was repeated daily for 8 consecutive days with *in vivo* calcium imaging and behavior recordings conducted during the cohousing period that followed defeat days 2, 5, and 8 (Def2, 5, 8). On day 9, the social interaction test (SIT) was conducted along with *in vivo* calcium imaging recording. (**b**) Schematic demonstrating the relative head distance (Dist_head_) and head angle (Angle_head_) between the subject mouse (black) and its cohoused neighbour (white). (**c**) Average Dist_head_ (*left*) and Angle_head_ (*right*) following defeat sessions 2, 5, and 8 in control (con; white), resilient (res; blue), and susceptible (sus; red) mice. (**d**) *Left:* Schematic of session 1 (empty enclosure) and 2 (CD1 present in enclosure) of the SIT along with representative trace of subject trajectory (blue). *Middle:* Dist_head_ during sessions 1 and 2 of the SIT. *Right:* Angle_head_ during sessions 1 and 2 of the SIT. All data are expressed in mean ± SEM. *p < 0.05, effect test of group with two-way ANOVA. †p < 0.05, ††p < 0.01, paired t-tests.

We compared the head distance (Dist_head_) and the angle of heading direction (Angle_head_) of subject mice to the neighboring social target (head) during cohousing, or the mouse-containing cup (center) in the SIT (**Fig. 1b**). During cohousing, resilient mice maintained a shorter Dist_head_ and a smaller Angle_head_ with aggressors compared to controls (**Fig. 1c**), suggesting closer contact between resilient mice and aggressors during cohousing. In addition, comparing the Dist_head_ and Angle_head_ during the SIT revealed a smaller Dist_head_ in resilient mice when compared to susceptible mice (**Fig. 1d**). Notably, only control and resilient mice, but not susceptible mice, displayed a between-session reduction of Dist_head_ in the SIT. The increase in Dist_head_ does not represent a lack of attention towards the aggressor by susceptible mice. Unlike control mice, both susceptible and resilient mice displayed a between-session reduction of Angle_head_, suggesting a smaller viewing angle of stressed mice toward the social target in the SIT. In addition, two-way ANOVA revealed a group effect of smaller Angle_head_ in susceptible mice when compared to resilient mice. Thus, during the SIT resilient mice remained at closer proximity and positioned towards the CD1, while susceptible mice maintained a further distance but continued to face the CD1, potentially indicators of vigilance or social anxiety (*25*).

To specifically analyze social behavior during cohousing, interactions were defined as behavioral frames where the Dist_head_ was <10 cm (D10) and the Angle_head_ between mice was <50° (A50; **Fig. 2a**). Both stressed groups spent more time at D10 than controls. We also compared the percentage of time when mice were situated at regions away from the social targets (Dfar, when Dist_head_ >25 cm) and did not find differences between the 3 mouse groups (**fig. S2a**). Next, we compared the percentage of time when Angle_head_ between mice was <50° (A50) and found that susceptible mice remained at A50 longer than controls (**Fig. 2a**). A significant increase in time at A50 was also seen in susceptible mice between the 2^nd^ and 8^th^ defeat session. Notably, when we compared the compass angle of subject mouse head direction (i.e., Angle_head_ towards north = 0, unrelated to social targets), we observed no between-group differences **(fig. S2b)**, suggesting the increased time spent at A50 in susceptible mice is social target-related. Finally, while both stressed groups moved less than control mice during recording, no difference between resilient and susceptible groups was found (**fig. S2c**). Together, nuances in social behavior appear to manifest throughout the course of defeat and result in diverging behavioral phenotypes in the SIT. These findings prompted us to further examine neuronal correlates of interactions (D10 and A50) between subject mice and social targets.

**Figure 2.**
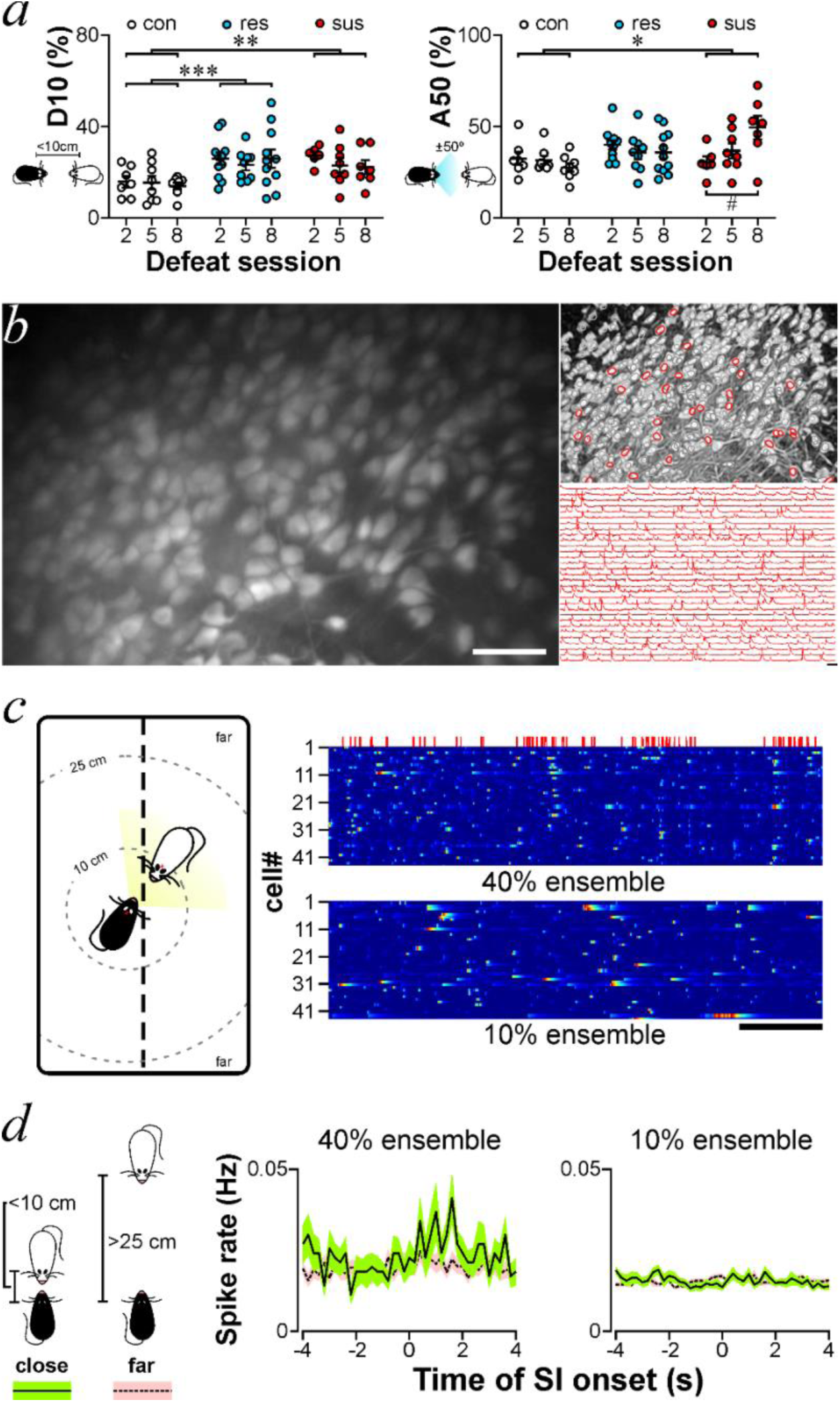
Identifying dCA1 social ensembles. (**a**) Percentage of time spent at (*left*) Dist_head_ < 10 cm (D10) and (*right*) Angle_head_ ±50 degrees (A50) following defeat sessions 2, 5, and 8 in control (con; white), resilient (res; blue), and susceptible (sus; red) mice. |(**b**) *Left:* dCA1 field of view demonstrating maximal activity from all active neurons expressing GCaMP6f in a recording session (scale bar = 20 μm). *Right:* Micrograph from *in vivo* calcium imaging recording outlining individual dCA1 neurons with traces representing corresponding GCaMP6f fluorescent signals (scale bar = 1 min). (**c**) *Left:* Schematic representing interactions between CSDS subject mouse (black) and CD1 aggressor (white) in the cohousing post-CSDS. *Right:* Inferred calcium activity for each dCA1 neuron sorted according to activity during social interaction bouts (red lines above raster plots). dCA1 social ensembles of neurons that were active during 40% (*top*) or 10% (*bottom*) of social interaction bouts (scale bar = 2 min). (**d**) Inferred calcium spike rate of (*left*) 40% ensemble and (*right*) 10% ensemble aligned according to close (< 10 cm, green) and far (> 25 cm, pink) social interaction onset. (**e**) Inferred calcium spike rate of social ensembles during close and far social interactions. All data are expressed in mean ± SEM. *p < 0.05, **p < 0.01, ***p < 0.001, two-way ANOVA followed by Tukey’s multiple comparisons test. †p < 0.05, ††p < 0.01, †††p < 0.001, paired t-tests.

### dCA1 social ensemble activity throughout CSDS and SIT

We next determined whether variations in social behavior were accompanied by changes in hippocampal activity as measured by *in vivo* calcium imaging. Fluorescent calcium spikes were extracted and used as a proxy for neuron activation (**Fig. 2b**). We found no between-group difference in inferred spike rate of dCA1 neurons across different defeat days (**fig. S2d**). These findings prompted us to identify dCA1 neurons that are active during social interaction. We extracted dCA1 population activity that is related to social interaction between the subject and target mice. This was done through the calculation of cosine similarity indexes (CSI; (*26*)) between the mean inferred calcium spiking rate vector from an ensemble of neurons and the behavioral vector of social interaction bouts (D10 and A50). We identified ensembles of neurons that were active in >40% of all social interaction bouts. We compared the activity of neuronal ensembles that were active in 40% vs. 10% of all social interaction bouts following a given defeat episode (**Fig. 2c**). An increase in inferred calcium spike rate of the 40% ensemble was observed at the onset of close interactions (Dist_head_ < 10 cm), compared to far encounters (Dist_head_ > 25 cm) (**Fig. 2d**). Inferred calcium spike rate of the 10% ensemble remained similar regardless of distance to the aggressor. These results indicated that the 40% ensemble reflected close social interactions with a greater fidelity than the 10% ensemble. For all future analyses, the 40% ensemble was used and will now be termed the social ensemble.

Across different sessions of CSDS cohousing, inferred spiking rate of social ensemble neurons from all groups was greater when subjects were near social targets compared to far, except in susceptible mice after the 2^nd^ session of social defeat (**Fig. 3a**). Although transient, the absence of social ensemble activity favoring close interactions in susceptibility indicated that the neuronal representation of social interactions may be diminished in susceptibility. We next quantified the level of congruence between their mean activity and social interactions that is represented by CSI. Social ensembles showed greater CSI in resilient mice compared to both susceptible and control mice (**Fig. 3b**), indicating that activity from social ensembles are better representative of interactions in resilience. Across CSDS, social ensemble size, which represents the percentage of recorded neurons in social ensembles, was also greater in resilient than control mice (**Fig. 3c**). Considering that neuron reactivation is another signature relating activity to an experience (*27*), we used cell registration (*23*) to track the activity of the same individual neurons across different social defeat sessions. On average, approximately 50% of neurons reactivate across CSDS cohousing (**Fig. 3d**). Individual neuron reactivation may be initiated by several cues including contextual features; thus, we focused on the reactivation of social ensemble neurons. Approximately 50-70% of social ensemble neurons reactivated across CSDS cohousing. To more specifically evaluate the activity of cells related to social interaction we isolated the neurons that continued to comprise the social ensemble across different days. We found that more social ensemble neurons continued to comprise the social ensemble at later timepoints in resilient mice compared to controls. Next, we compared the reactivation of dCA1 neurons recorded in CSDS cohousing during SIT (**Fig. 3e**). We found that susceptible mice displayed a reduction of reactivation of social ensemble neurons than resilient and control mice. These findings suggest that in resilient mice, social ensemble neurons more consistently represent interaction across different defeat days.

**Figure 3.**
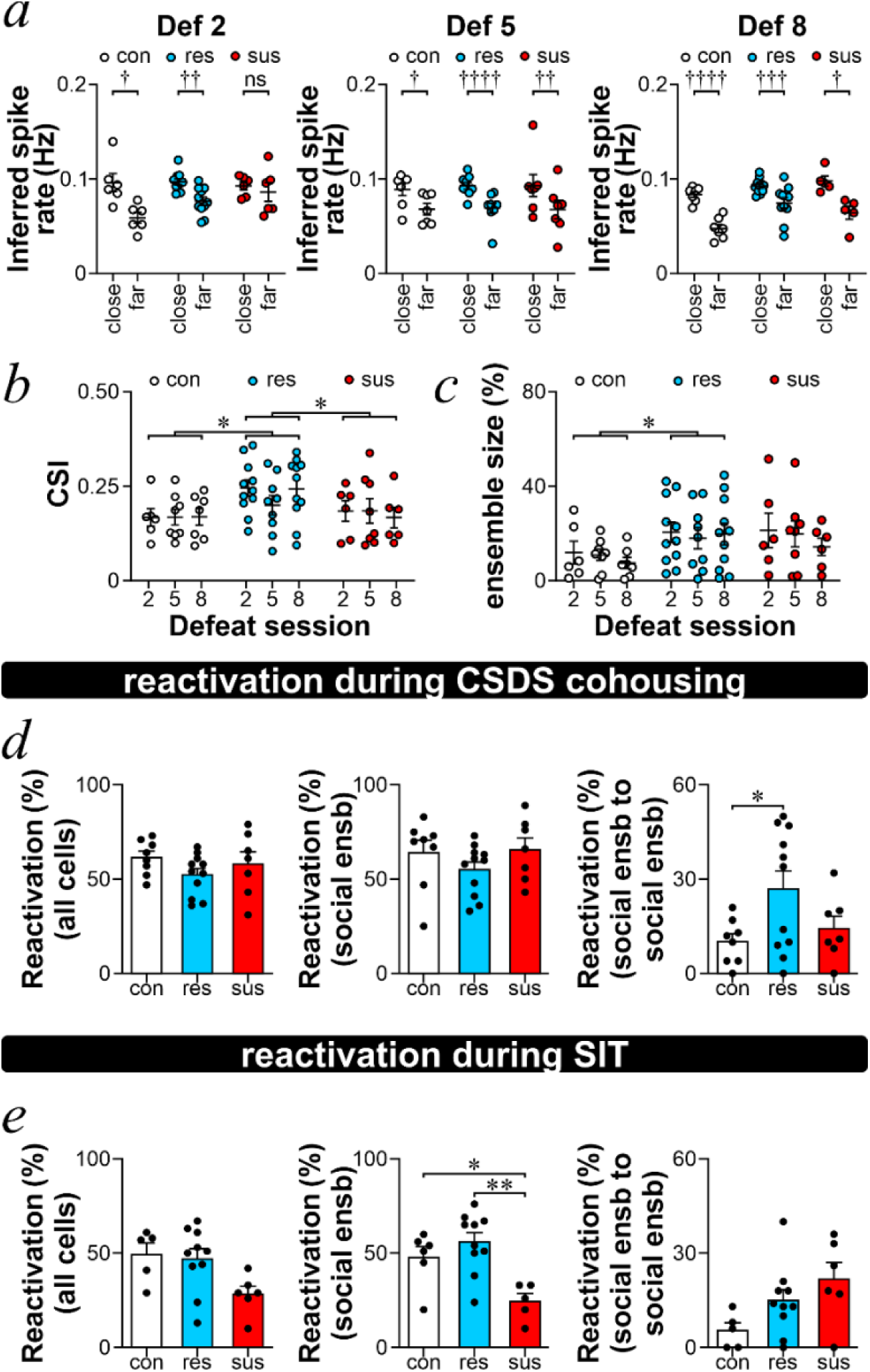
dCA1 social ensembles better represent interactions throughout CSDS in resilient mice. **(b)** Average cosine similarity index (CSI) of social ensemble neurons following defeat sessions 2, 5, and 8 in control (con; white), resilient (res; blue), and susceptible (sus; red) mice. *p < 0.05, effect test of group with two-way ANOVA. (**c**) Social ensemble size. *p < 0.05, effect test of group with two-way ANOVA. (**d**) Average percentage of reactivated dCA1 neurons in defeat sessions 2, 5, and 8. Histograms represent reactivations of (*left*) all recorded cells; (*middle*) social ensemble neurons; (*right*) social ensemble neurons that continue to comprise the social ensemble. All data are expressed in mean ± SEM. *p < 0.05, One-way ANOVA followed by Tukey’s multiple comparisons test. (**e**) Average percentage of reactivated dCA1 neurons in the social interaction test (SIT). Histograms represent reactivations of (*left*) all recorded cells; (*middle*) social ensemble neurons; (*right*) social ensemble neurons that continue to comprise the social ensemble. All data are expressed in mean ± SEM. *p < 0.05, **p < 0.01, One-way ANOVA followed by Tukey’s multiple comparisons test.

To determine whether variations in dCA1 social ensemble activity during CSDS cohousing and SIT can be attributed to parallel differences in interactions we compared behavior in these recording sessions. The number of social bouts during CSDS cohousing was similar between the 3 mouse groups (**Fig. 4a**). In those social bouts, both defeated groups initiated approach less frequently and reacted to interactions with escape behavior more than control mice (**Fig. 4b**). As expected, susceptible mice interacted less with the aggressors in the SIT. Together, observed differences in social ensemble activity during CSDS cohousing and SIT are likely not explained by the dynamics of social behavior. Lastly, it is possible that dCA1 place cell properties are responsible for the differences in ensemble activity we observed in different mouse groups. We found that lower mean spatial information was carried by the social ensemble in both defeated groups compared to control mice during CSDS (**Fig. 4c**). These findings seem to be congruent to the reported detrimental effect of stress on place cell properties (*28*). Notably, social ensemble activity during cohousing is likely less sensitive to changes in place cell properties since all mice during cohousing were freely moving and interacting at random locations.

**Figure 4.**
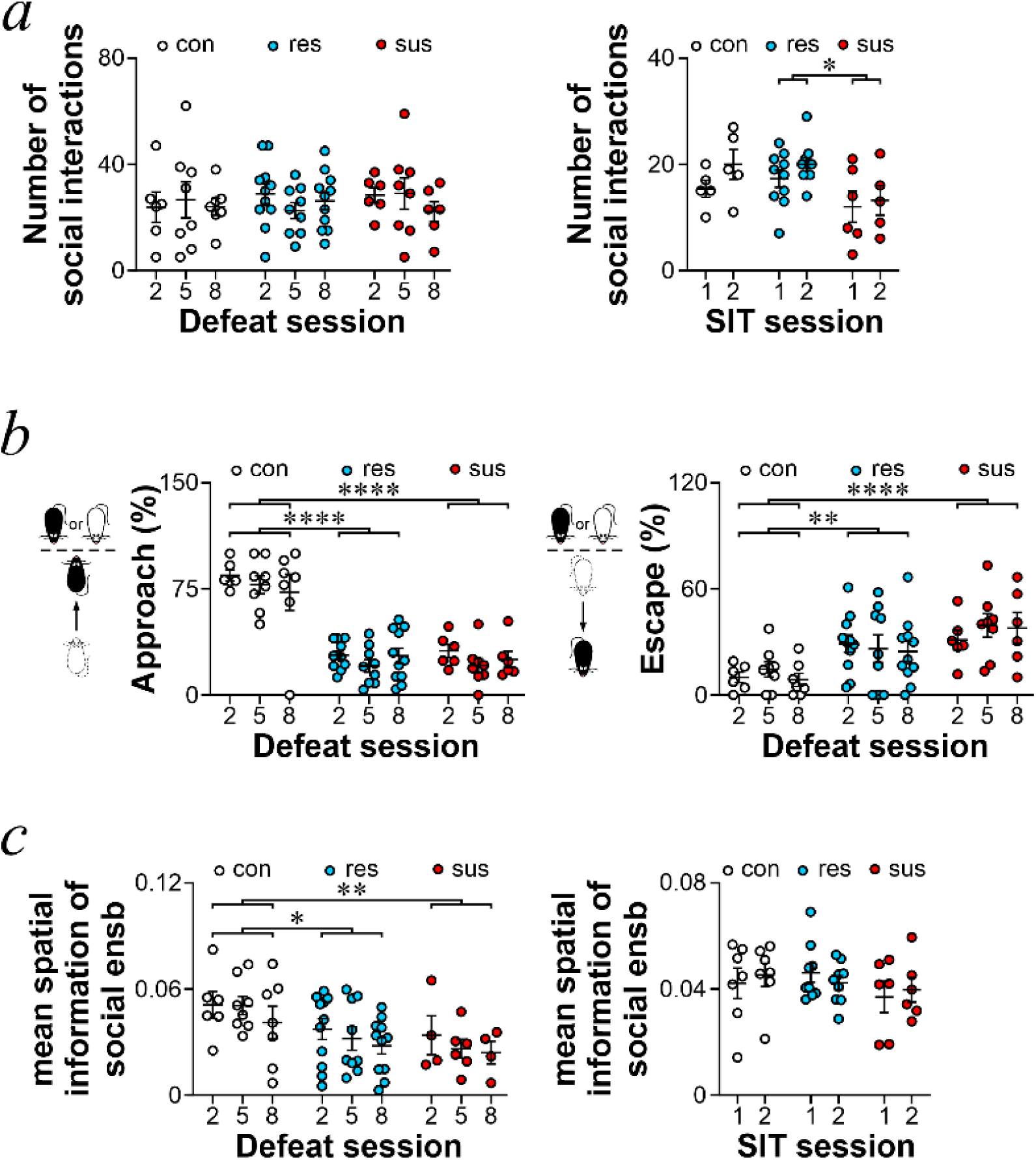
Social behavior and activity governed by spatial information do not account for decreased social ensemble fidelity in CSDS susceptibility. (**a**) Number of social interactions (Dist_head_ < 10 cm and Angle_head_ ± 50°) with CD1 aggressors or same-strain controls following defeat sessions 2, 5, and 8 in control (con; white), resilient (res; blue), and susceptible (sus; red) mice. (**b**) Percentage of (*left*) approaches and (*right*) escapes initiated by C57BL/6 subject mice during CSDS cohousing with CD1 aggressors or same-strain controls. (**c**) Mean spatial information represented by dCA1 neurons in the social ensemble. All data are expressed in mean ± SEM. *p < 0.05, **p < 0.01, ****p < 0.0001, effect test of group with two-way ANOVA.

### CSDS impairs social memory with a greater effect in susceptible mice

The decreases in social ensemble fidelity in CSDS susceptibility may result in deficits in social memory. As shown by previous studies and our findings from another cohort of mice **(fig. S3 & S4)**, avoidance in CSDS susceptible mice is specific to the CD1 strain, suggesting a contribution from social memory for this aggressive strain. Using the preference for mice to interact with a novel conspecific (*29*), we characterized the impact of CSDS on social memory for the CD1 strain. We studied 3 different measures of social memory 4 days post-SIT (**Fig. 5a**): *short-term social memory* for a familiar CD1 measured 2 hours later; *social novelty memory* for a novel CD1 vs. a familiar CD1; and *long-term social memory* for a familiar CD1 measured 1 day later. In all cases, control mice displayed a decrease in interaction time when the social target was less novel (**Fig. 5b**). Stressed mice showed an impairment in social memory across all 3 measures when compared to controls. Interestingly, susceptible mice exhibited lower habituation to the revisited familiar CD1 mouse and lower preference towards the novel CD1 mouse when compared to resilient mice. Therefore, CSDS impairs social memory for the aggressor strain, but the effect is greater in susceptible mice.

**Figure 5.**
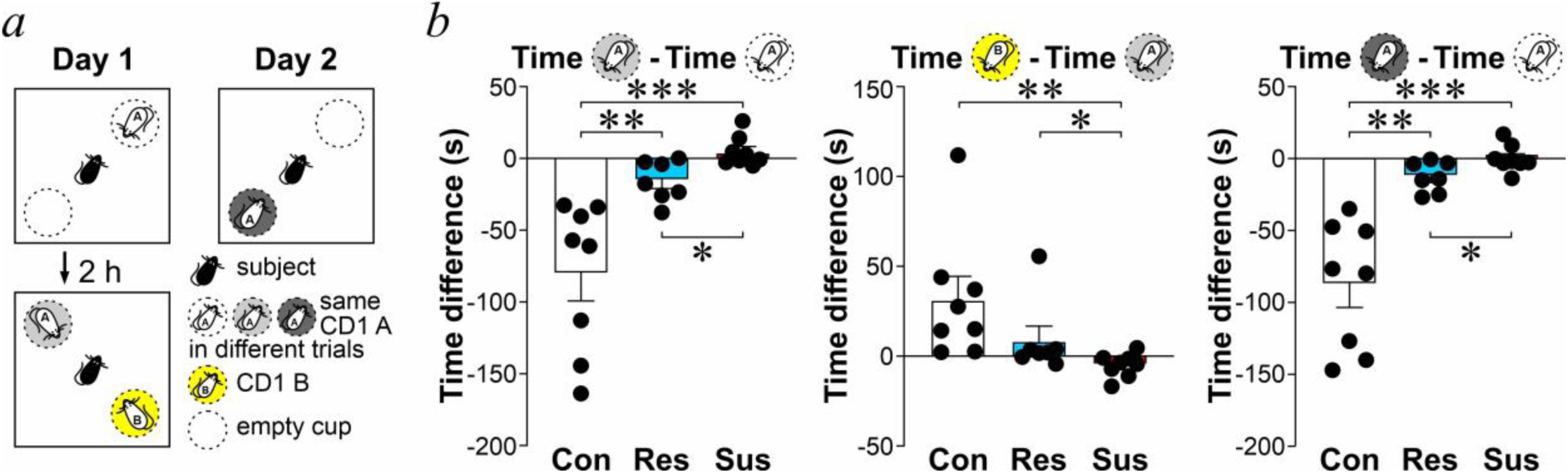
CSDS impairs social memory to a greater extent in susceptible mice. **(a)** Schematic of social memory tests. On day 1, subjects (defeated mice or non-stressed controls) interacted with novel CD1 A (white) and an empty cup in an open field. Two hours later, subjects could investigate the now familiar CD1 A (light grey) or the novel CD1 B (yellow). On day 2, subjects could investigate familiar CD1 A (dark grey) or an empty cup. **(b)** Social memory performance in control (con; white; n = 8)), resilient (res; blue; n = 7), and susceptible (sus; red; n = 8) mice. *Left:* Short-term social memory measured as the time spent investigating CD1 A when it was familiar (2h) compared to novel. *Middle:* Social novelty preference measured as the time spent investigating novel CD1 B compared to familiar CD1 A. *Right:* Long-term social memory measured as the time spent investigating CD1 A when it was familiar (24 h) compared to novel. All data are expressed in mean ± SEM. *p < 0.05, **p < 0.01, ***p < 0.001, Kruskal-Wallis test followed by Wilcoxon test.

## Discussion

Using *in vivo* calcium imaging to longitudinally track hippocampal activity throughout CSDS, we identified differences in dCA1 representations of social interactions between susceptible and resilient groups. In examining population activity, we found that both the size and the information carried by social ensembles during CSDS cohousing were higher in resilient mice. Additionally, susceptible mice showed decreased social representation and weaker social memory for CD1 mice. To our knowledge, this is the first report of the evolution of dCA1 activity throughout CSDS. Our findings suggest that differences in the processing of social information by the dCA1 could regulate stress susceptibility.

CSDS is a commonly used rodent model for the study of diverging responses to social stressors (*7*). Following identical stressors, the SIT can separate those that are susceptible and expressing social avoidance from resilient mice that retain a greater level of social investigation (**fig. S1 and S3**). Both susceptible and resilient mice shared behavioral similarities during CSDS cohousing. Compared to control mice, susceptible and resilient mice were less active to initiate social interactions with their cohoused social target and initiated escapes more frequently (**Fig. 4**). Both defeated groups also traveled less than control mice but spent more time at a closer head distance to their cohoused social target (**fig. S2, Fig. 2**). These findings indicate that both susceptible and resilient mice appeared vigilant towards their neighboring aggressor during cohousing. However, subtle differences were also found. Resilient mice maintained a shorter Dist_head_ and Angle_head_ to the CD1 aggressor than control mice during cohousing (**Fig. 1**). Comparing the occupancy of A50, only susceptible mice showed a smaller Angle_head_ to cohoused CD1 aggressors compared to controls, particularly following the last defeat episode (**Fig. 2**). These findings support the importance of considering both head distance and heading direction towards social targets in our definition of social ensembles.

The dCA1 is commonly known for its role in spatial navigation. Notably, dCA1 spatial representations can be modulated by social processes and valenced experiences. dCA1 social place cells representing the location of social targets could support navigation processes through social environments (*13, 14*). Recently, dCA1 activity in bats was shown to simultaneously represent spatial and social information (*30*). Cells tuned to the relative distance between a subject and another conspecific were identified in instances where social presence was behaviourally important, such as the avoidance of collisions during bat flight (*31*). Spatial mapping is also influenced by valenced experiences, such that dCA1 place cells are biased to represent locations associated with reward (*32–34*) and remap following an aversive experience (*35*). dCA1 neuron ensembles tuned to aversive stimuli have also been identified (*36*). Finally, dCA1 activity is required for identifying social targets that were associated with emotionally salient experiences (*15, 16*). Here, we additionally uncover differences in the dCA1 representation of social interactions in CSDS susceptible and resilient mice.

Compared to anatomical methods labelling active neurons (*8, 37*), *in vivo* calcium imaging allows for increased temporal reliability that enabled us to isolate neurons whose activity covaried with behavior. How these changes in dCA1 function contribute to CSDS susceptibility and resilience remains unclear. Following the 2^nd^ episode of CSDS, activity of dCA1 social ensembles was reduced at a far distance from the social target in control and resilient mice (**Fig. 3**), supporting a reduction in social ensemble selectivity for close social interactions in susceptibility. Susceptible mice also exhibited lower reactivation of social ensembles in the SIT (**Fig. 3**). These findings point to deficits in social information processing and memory in susceptible mice, which may indicate a vulnerability to social stress impacting cognitive function. In fact, impairments in spatial cognition have been reported following CSDS (*38, 39*) and are accompanied by decreased hippocampal levels of proteins supporting memory functions (*40*). Alternatively, the deficits in the dCA1 representation of social interaction could reduce the quality of social memory in susceptibility, supporting memory generalization processes. CSDS has been shown to induce changes in dCA1 activity and contextual fear generalization (*41*). Despite both susceptible and resilient mice differentiating between CD1 and C57BL/6 mice in the SIT (**fig. S4**), susceptible mice showed greater difficulty recognizing novel vs. familiar CD1 mice in the social memory test (**Fig. 5**). Moreover, hippocampal corticosterone, which is elevated following just a single social defeat episode (*42*), could facilitate fear generalization (*43*).

Alternatively, resilience may be reflective of a better representation of social and environmental cues. Social ensembles had increased CSI, greater size, and reactivated more frequently across different CSDS episodes in resilient mice only (**Fig. 3**). Sociability is preserved in CSDS resilient mice interacting with same strain adults or juveniles, potentially attributed to the recognition of their non-threatening nature (*44, 45*). Similarly, resilient mice may be recognizing that the social target never previously attacked them or that there is a lack of direct physical contact with the confined aggressor signifying it is not an immediate threat.

We may draw parallels to hallmark memory deficits observed clinically in stress-related psychopathologies. We previously utilized an activity-dependent immunohistochemical technique and observed increased reactivation of dCA1 neurons active during CSDS in susceptible mice (*8*). This may be reflective of a negative memory bias, or a greater memory for negative events, which is the most robust cognitive symptom of depression (*46*). Unlike examining the effect of the defeat experience on dCA1 activity, our current study used *in vivo* calcium imaging to measure dCA1 activity relating to social behavior throughout the course of CSDS. Using a technique with greater temporal resolution revealed more nuanced stress-induced changes, suggesting imprecise memory content in susceptibility. In fact, poor recollection of detailed autobiographical memory is a feature of depression and post-traumatic stress disorder (*47*). Similarly, fear generalization, or defensive responses triggered by incorrect assessments of imminent threat, is well-described in post-traumatic stress disorder and anxiety (*48*). Taken together, it is possible that susceptible mice have increased memory for the negative CSDS event, but that this memory is lacking specificity.

Importantly, investigations in female cohorts are warranted as stress-related psychopathology is more prevalent among women (*49, 50*) presenting a limitation of the current study. CSDS models have typically been more challenging to implement in female cohorts due to a lack of consistent natural aggressive behavior among female mice. With the advent of promising new defeat models, future studies related to social memory and information processing following social stress can be conducted in both sexes (*51*).

In sum, our findings uncover a role for hippocampal social cognitive processes in the individual variability of stress responses. CSDS resilience is characterized by greater social investigation of the aggressor strain and is accompanied by more reliable dCA1 correlates of social interaction and maintained social memory, while these changes are absent in susceptible mice who instead show diminished social memory for the aggressor strain. Susceptibility may then manifest in mice through deficits in the processing and storage of social information.

## Acknowledgements

The present study used the services of the Molecular and Cellular Microscopy Platform in the Douglas Hospital Research Centre. Melina Jaramillo Garcia helped set up the imaging verification.

## Author contributions

Conceptualization: AL, TPW

Methodology: AL, TRZ, ASW, XLYL, CYHF, BCMF, TPW

Investigation: AL, TRZ, ASW, XLYL, TPW

Funding acquisition: TPW

Project administration: TPW

Supervision: TPW

Writing – original draft: AL, TPW

Writing – review & editing: AL, TPW

## Data and materials availability

All data are available in the main text or the supplementary materials. Data and codes are available upon request.

