## Supplementary Material for "Diminished social memory and hippocampal correlates of social interactions in chronic social defeat stress susceptibility"

**Supplementary figures**

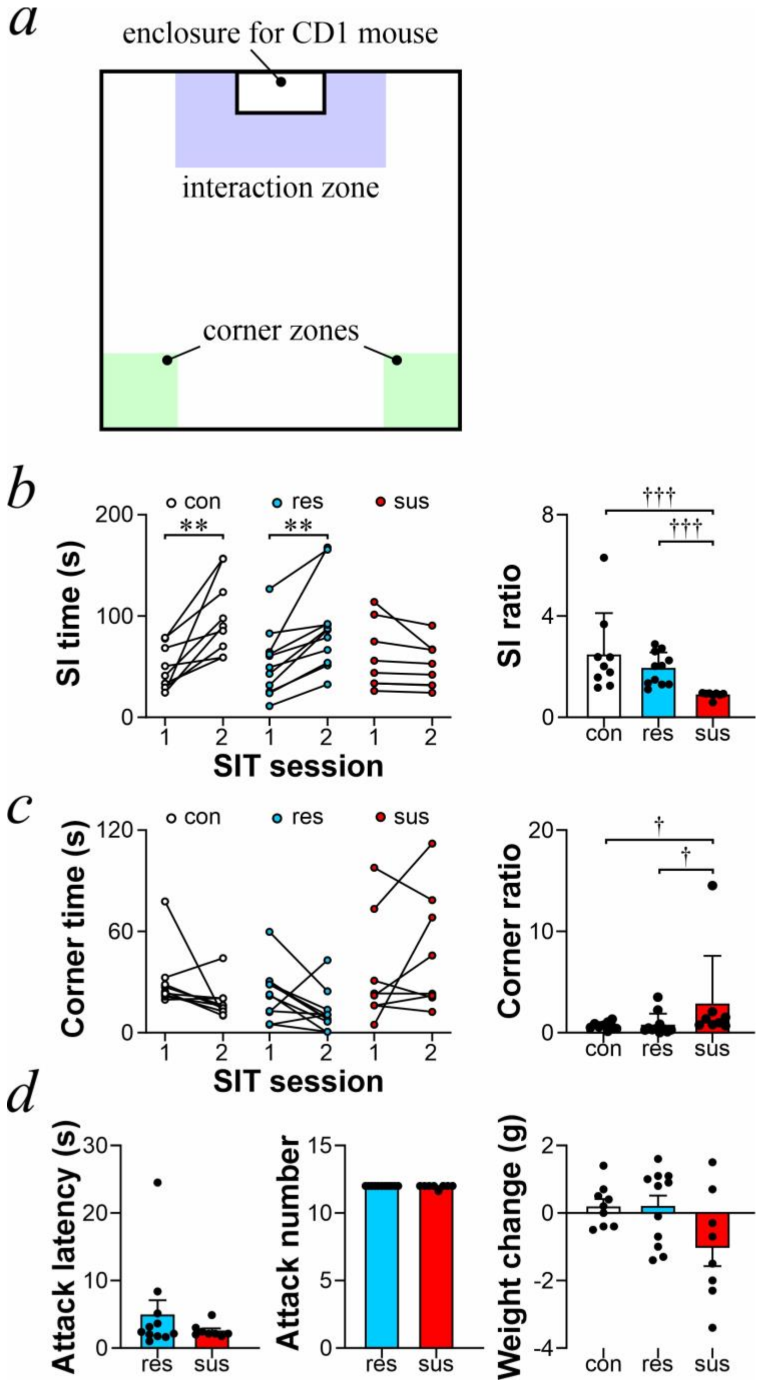


**Supplementary Figure 1. Susceptible mice outfitted for *in vivo* calcium imaging show social avoidance in SIT despite similar CSDS experience.**

**(a)** A schematic diagram of the open field in the social interaction test (SIT). The virtual interaction zone (purple) and corner zones (green) are indicated.

**(b)** *Left:* Social interaction time in session 1 (empty enclosure) and session 2 (CD1 present in enclosure) of the SIT in control (con; white; n = 9), resilient (res; blue; n = 11), and susceptible (sus; red; n = 8) mice. *Right:* Social interaction ratio (time spent in interaction zone during session 2/time spent in interaction zone during session 1).

**(c)** *Left:* Corner time in session 1 and session 2 of the SIT. *Right:* Corner ratio (time spent in corner zones during session 2/time spent in corner zones during session 1).

**(d)** Mean attack latency, attack number and weight change throughout CSDS. All data are expressed in mean ± SEM, $^{**}p < 0.01$, paired t-test. $^{\dagger}p < 0.05$, $^{\dagger\dagger\dagger}p < 0.001$ Kruskal-Wallis test followed by Wilcoxon test.

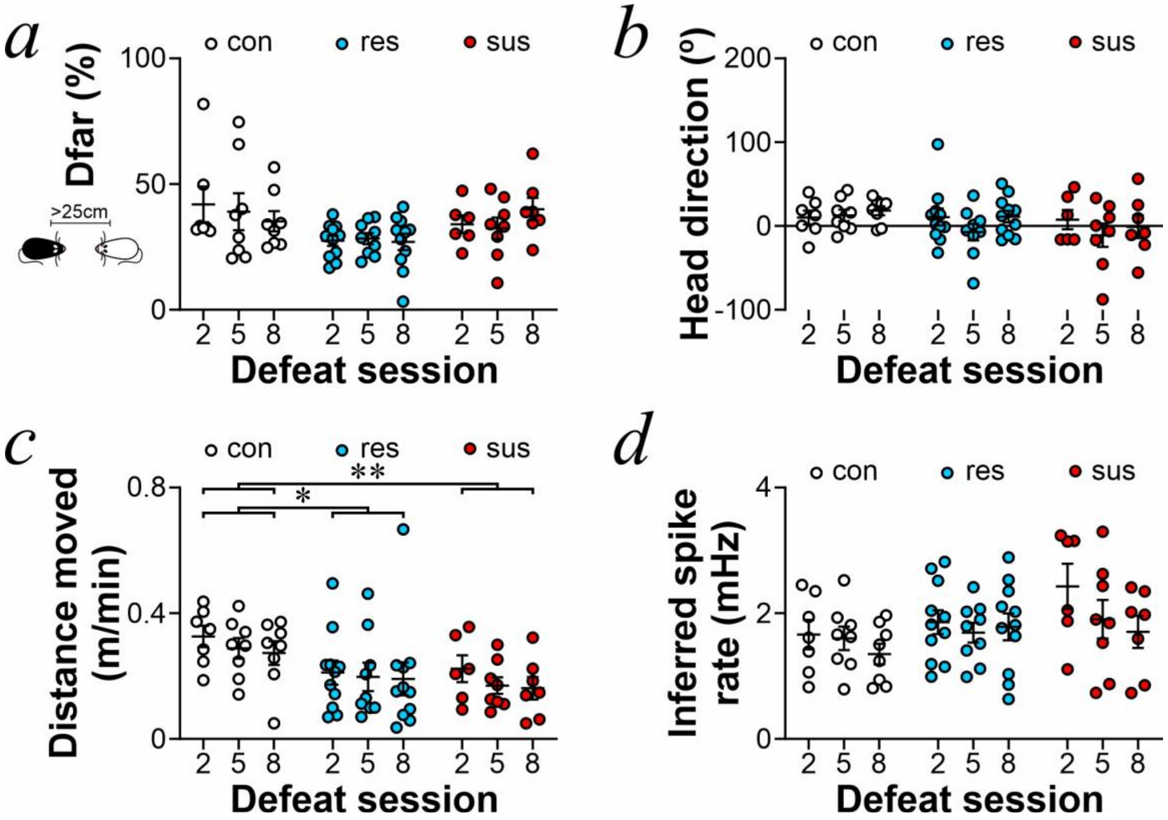


**Supplementary Figure 2. Measures of behavior and dCA1 inferred spiking rate during CSDS cohousing.**

**(a)** Percentage of time spent at $Dist_{head}$ > 25 cm (Dfar) following defeat sessions 2, 5, and 8 in control (con; white), resilient (res; blue), and susceptible (sus; red) mice.

**(b)** Average head direction.

**(c)** Average distance moved (m/min).

**(d)** Inferred spike rate (mHz) of all recorded dCA1 neurons in a session. All data are expressed in mean ± SEM. $^*p < 0.05$, $^{**}p < 0.01$, effect test of group with two-way ANOVA.

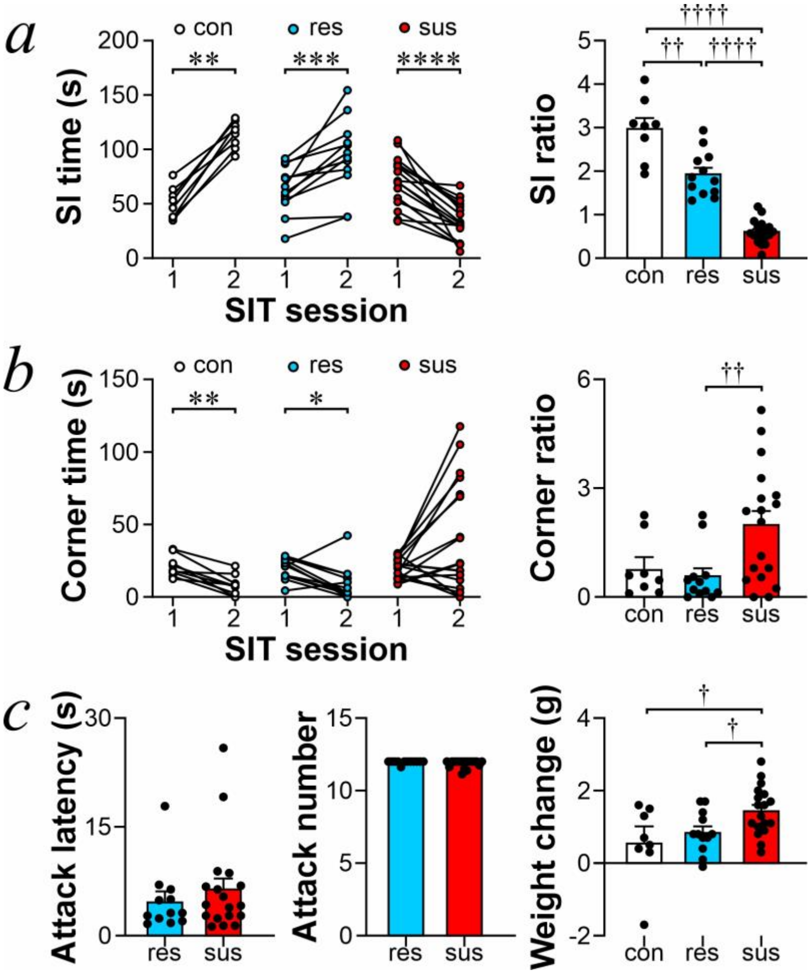


**Supplementary Figure 3. Susceptible mice show social avoidance in SIT despite similar CSDS experience.**

**(a)** *Left:* Social interaction time in session 1 (empty enclosure) and session 2 (CD1 present in enclosure) of the SIT in control (con; white; n = 8), resilient (res; blue; n = 12), and susceptible (sus; red; n = 18) mice. *Right:* Social interaction ratio (time spent in interaction zone during session 2/time spent in interaction zone during session 1).

**(b)** *Left:* Corner time in session 1 and session 2 of the SIT. *Right:* Corner ratio (time spent in corner zones during session 2/time spent in corner zones during session 1).

**(c)** Mean attack latency, attack number and weight change throughout CSDS. All data are expressed in mean ± SEM. *$p < 0.05$, **$p < 0.01$, ***$p < 0.001$, ****$p < 0.0001$, paired t-test. †$p < 0.05$, ††$p < 0.01$, ††††$p < 0.0001$, Kruskal-Wallis test followed by Wilcoxon test.

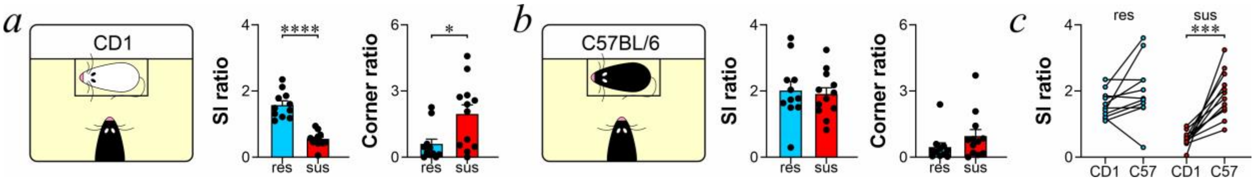


**Supplementary Figure 4. CSDS susceptibility does not remain stable after retesting with C57BL/6 mouse in the SIT**

Social interaction ratio (time spent in interaction zone during session 2/time spent in interaction zone during session 1) and corner ratio (time spent in corner zones during session 2/time spent in corner zones during session 1) during the SIT using **(a)** a CD1 or **(b)** C57BL/6 social target in resilient (res; blue; n = 11) and susceptible (sus; red; n = 12) mice.

(c) SI ratio when tested with CD1 and C57BL/6 social targets. All data are expressed in mean ± SEM. *p < 0.05, ****p < 0.0001, Student's t-test, non-paired in (a) and (b), paired in (c).
